# A high-resolution, sex-stratified atlas of transcriptional and alternative splicing dynamics in the fasting mouse liver

**DOI:** 10.64898/2026.04.28.720569

**Authors:** Jess Treeby, Shin-Yu Kung, Ben Saer, Judith Gramunt Prillo, Ann Louise Hunter, Jean-Michel Fustin

## Abstract

The liver’s response to fasting is a fundamental adaptive mechanism, yet its temporal complexity, circadian dimension and sexual dimorphism remain incompletely understood. Here we present a high-resolution transcriptomic atlas of the fasting mouse liver, profiling both male and female C57BL/6 mice every 4 hours across a 36-hour fast initiated at a defined circadian time.

Using an integrated framework combining high-resolution time-series analysis with stringent statistical filtering, we identify 2,995 genes organised into eight kinetically distinct clusters that collectively delineate a phased transcriptional transition from acute metabolic remodelling to prolonged nutrient deprivation. Contrary to prior reports, fasting amplifies rather than diminishes circadian transcription, increasing the number of rhythmically expressed genes from 727 to 1,233—indicative of a clock re-gearing rather than clock disruption.

Extending our analysis to exon-level quantification reveals that differential exon usage constitutes an independent regulatory layer, targeting processes not captured by transcript abundance alone, including SAM metabolism, ferroptosis and mRNA processing. Circadian rhythmicity in exon usage is, to our knowledge, demonstrated here for the first time. Sexual dimorphism is pervasive but primarily quantitative, reflecting differences in the magnitude and temporal precision of a conserved programme rather than divergent regulatory logic. This open-access, multi-layered dataset provides a comprehensive resource for the study of nutritional and circadian regulation of liver metabolism.

## INTRODUCTION

The hepatic response to fasting is a fundamental survival mechanism, yet our understanding of its temporal and molecular complexity remains fragmented.

Early transcriptomic efforts with microarrays identified only a few hundred genes from a pool of 20,000 probes^1^, with fasting durations of 12, 24 and 72 hours and limited clarity regarding the exact onset of food withdrawal. Other studies established important links between food availability and health^2–4^, but these utilised subtraction microarrays or restricted time points that could not capture the fluid kinetic transitions of the fasting liver.

A critical component of the response to fasting is the integration of circadian timing with nutritional status. While it is known that a 24-hour fast initiated during the day versus at night produces distinct hepatic transcriptome signatures^5^, many studies do not standardise the starting *Zeitgeber* or Circadian Time (ZT or CT, with ZT0 or ZT12 respectively defined as the start of the light or dark phase and CT0 or CT12 as the start of the rest or active phase in mice housed in constant darkness). Since mice consume most of their food during their active phase at night^6,7^, a fast of identical duration started in the morning versus the evening will have dramatically different effects on liver physiology that are not necessarily dependent on the endogenous circadian system.

Studies focusing on circadian rhythms of gene expression present distinct methodological limitations that obscure the adaptive kinetics of the fasting liver^8,9^. In one study^8^, mice were subjected to a ~24-hour fast starting at either CT04 or CT16, and gene expression was profiled the following day every 2 hours; at the earliest sampling point, animals had already been fasted for 20–22 hours, capturing prolonged starvation rather than the dynamic fed-to-fasted transition. Another study^9^ sampled livers after a fixed 24-hour fast initiated at six different times of day, creating a static snapshot that cannot resolve the immediate post-withdrawal response. Both studies reported a substantial decrease in rhythmic gene number, concluding that feeding–fasting cycles, rather than the endogenous circadian clock, are the primary driver of rhythmic hepatic transcription.

Perhaps the most significant gap in the field is the lack of sex-stratified data. Although Della Torre et al. (2018)^10^ identified amino acid metabolism as a major sex-discriminating factor during short-term fasting, much of the foundational work on fasting-regulated transcriptional pathways has relied exclusively on male cohorts. A high-resolution, longitudinal dataset tracking both sexes simultaneously—to determine whether metabolic adaptation follows a shared blueprint or divergent strategies—is currently lacking.

Our understanding of the fasting response has also been largely restricted to bulk transcript abundance, leaving the alternative transcriptome significantly under-explored. Differential exon usage allows a single gene to produce distinct protein isoforms or regulatory transcripts, yet evidence for fasting-induced splicing in the liver is limited to a small number of studies examining male mice fasted for 15 hours from an unspecified time^11^ or the exon usage of a narrow set of gene targets after a 14- or 24-hour fast from ZT15^12^. Identifying such shifts is crucial: alternative splicing can alter protein domains, induce nonsense-mediated decay, and thereby dictate cellular signalling and metabolic flux independently of total transcript levels.

### Transcriptional response to fasting

To address these gaps, we profiled the hepatic transcriptome of both male and female C57BL/6 mice following food withdrawal initiated at CT10—two hours before the onset of the dark/active phase (CT12). Liver tissue was harvested every 4 hours from *ad libitum*-fed and fasted animals across a continuous 36-hour window (CT08– CT44), providing the temporal resolution needed to detect both early (2–10 hours) and late (10–34 hours) transcriptional changes relative to a common baseline. To identify the most robustly regulated genes, we combined time-series trend analysis using the R package maSigPro^13,14^ with pairwise comparisons by DESeq2^15^, and make all raw data freely available.

Our integrated analysis identified 2,995 high-confidence genes meeting both criteria: (1) adjusted *p* value < 0.05 and |log2 fold-change| > 1 at a minimum of two time points by DESeq2 (Supplemental Table S1); and (2) significantly different expression trends between fasted and *ad libitum* livers by maSigPro (Figure 1A, Supplemental Table S2). These genes were organised into eight clusters that collectively delineate a phased transcriptional transition into the fasting state (Figure 1A, Supplemental Table S2). Clusters 1–4 and 8 characterise the acute transition phase, with genes undergoing rapid induction or repression within the first 2–10 hours of food deprivation (CT12–CT20) before reaching a new steady level. Clusters 5 and 7 display a protracted, progressive kinetic, with expression levels climbing steadily throughout the time course to peak after 24–32 hours of fasting, particularly in females.

**Figure 1:**
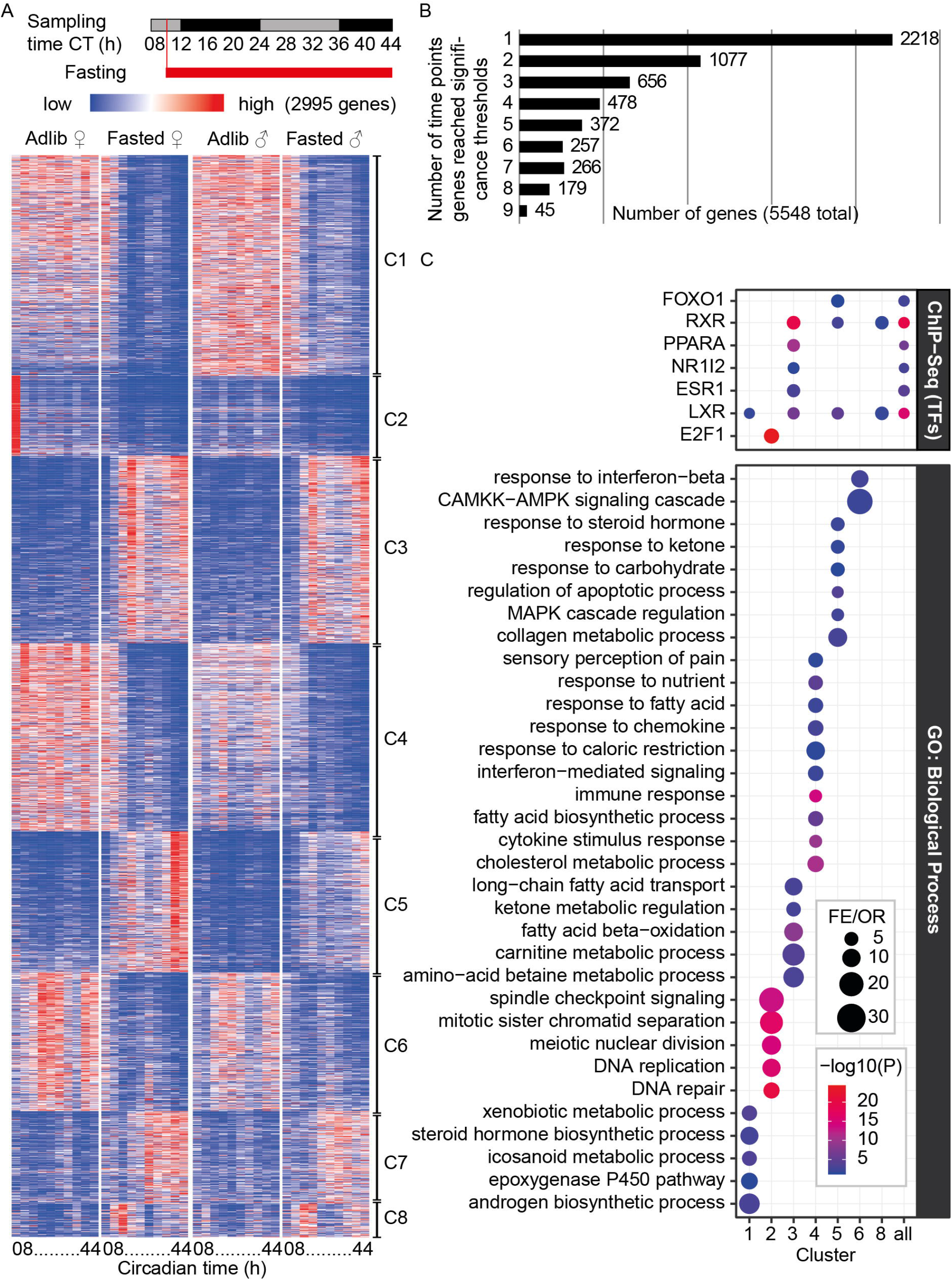
Phased transcriptional transitions and kinetic clustering of the hepatic fasting response. **A**, Heatmap of high-confidence genes passing all selection thresholds (adjusted *p* value < 0.05, |log2 fold-change| > 1 in at least two time points by DESeq2, and significant trend differences by maSigPro, *p* < 0.05), clustered by kinetic profile. Colours represent the mean of N = 2 males or 2 females; DESeq2 and maSigPro were performed with N = 4 mice (2♂ + 2♀). **B**, Frequency distribution of fasting-responsive genes by the number of time points at which DESeq2 significance thresholds were maintained. **C**, Top results from upstream regulatory network (Enrichr, top) and GO analyses (PANTHER, bottom) for each cluster in A. FE/OR = Fold Enrichment/Odds Ratio. See also Supplemental Table S1 and S2.

A distinct feature of the dataset is the interplay between directional fasting effects and underlying temporal oscillations. Clusters 1, 3, 5 and 8 exhibit a hybridised dynamic in which a near-linear progressive component is superimposed on a rhythmic oscillation. By contrast, the temporal behaviour of Cluster 6 is defined by attenuated rhythmicity: genes in this cluster display robust waveforms under *ad libitum* conditions that undergo progressive amplitude dampening as fasting continues—an observation that resonates with the reduced rhythmicity reported in prior studies^8,9^ and is revisited in the Discussion. These 2,995 genes represent 16.6% of the 18,023 genes retained after filtering for low counts, 3.8% of all annotated genes, and 89.9% of genes significant in at least two time points by DESeq2 pairwise comparisons (Figure 1B, Supplemental Table S1).

To elucidate the biological significance and regulatory architecture of these temporal profiles, we performed enrichment analysis on each cluster using PANTHER Gene Ontology (GO)^16,17^ and Enrichr (ChIP-X Enrichment Analysis)^18–20^ (Figure 1C, Supplemental Table S2).

The acute response phase (Clusters 1, 3, 4 and 8) was characterised by rapid metabolic remodelling. Cluster 3 emerged as a central metabolic hub, showing robust induction of fatty acid *β*-oxidation, ketone metabolism and carnitine transport, tightly coupled with enrichment of target genes for PPARA, RXR, LXR and NR1I2 (PXR)—signalling the activation of the hepatic lipid-catabolic engine (Figure 1C, Supplemental Table S2). In contrast, Cluster 2 was massively enriched for DNA replication and mitotic machinery, associated with E2F1 binding profiles, reflecting a programmatic suppression of cell proliferation and DNA synthesis (Figure 1C, Supplemental Table S2). No significant GO terms were obtained for Clusters 7 and 8.

The late-stage progressive response (Cluster 5) was defined by adaptation to prolonged nutrient deprivation, featuring pathways involved in ketone body metabolism and apoptotic regulation, with regulatory signatures for FOXO1, RXR and LXR (Figure 1C, Supplemental Table S2)—transcription factors that bridge the circadian clock and the metabolic fasting response^9,21,22^. Enrichment of ESR1 (Estrogen Receptor 1) target genes was identified in Cluster 3 and, to a lesser extent, Cluster 5 (q = 0.056), supporting a role for ESR1 in hepatic metabolic homeostasis in both sexes^23,24^ (Figure 1A, Supplemental Table S2).

Taken together, clusters characterised by net induction (Clusters 3, 5, 7 and 8) define a core catabolic programme driven by nuclear receptor signalling, prioritising mitochondrial import and *β*-oxidation of long-chain fatty acids to generate acetyl-CoA and ketone bodies. This is counterbalanced by the downregulation of Clusters 1, 2, 4 and 6, which orchestrate suppression of energy-intensive processes including cell proliferation and biosynthesis of fatty acids and steroid hormones (Figure 1C, Supplemental Table S2). Cluster 4 is notable for the breadth of its enrichment, spanning fatty acid and cholesterol synthesis, immune signalling and nutrient sensing, reflecting a broad, synchronised shutdown of hepatic biosynthetic capacity during fasting.

To identify sexual dimorphism in the transcriptional fasting response, we analysed males and females separately using maSigPro with the same DESeq2 thresholds. This identified 71 male-specific and 244 female-specific significant genes, alongside 2,735 shared genes (total 3,050; Figure S1A–C, Supplemental Table S1). The marginal number of genes detected exclusively by sex-stratified analysis indicates that sexual dimorphism in the fasting transcriptome is primarily quantitative—a difference in the magnitude of a shared response—rather than qualitative.

### Fasting amplifies circadian transcription

The non-linear expression dynamics observed in many clusters suggested that endogenous timing mechanisms compound the transcriptional effects of fasting. Prior work has shown that a fixed-duration fast initiated at different times of day reduces rhythmic gene number in the liver by ~33%^8,9^, and that restricted feeding can entrain the hepatic circadian clock^25^. In another experiment, oscillations of some clock and clock-controlled genes were shown to be affected during a fasting schedule that lasted for 24-46 hours^26^. More recently, *Per1* was found to be acutely induced by a 16-hour fast in the liver and was shown to be critical for the shift in fuel utilisation during nutrient deprivation^27^. However, no study has examined how the circadian transcriptome adapts from the very onset of fasting, or whether this adaptation is sexually dimorphic.

To address this, we analysed our data with dryR^28^, an R package that uses a Bayesian model selection framework to test for changes in circadian rhythmicity between conditions (Figure 2, Supplemental Table S3). Applying stringent dual criteria (DESeq2: q < 0.05, |log2FC| > 1 in ≥2 time points; dryR: BICW > 0.8), we identified 1,731 high-confidence genes: 132 lost rhythmicity in fasted livers (model 2), predominantly genes peaking during the rest phase (~CT7–8); 638 gained rhythmicity (model 3), distributed in two antiphasic groups peaking during the rest or active phase; 369 had altered phase or amplitude (model 5), mostly peaking during the active phase; 226 maintained identical rhythms, 204 of them with a shifted baseline (model 4); and 366 were arrhythmic in both conditions (Figure 2A–B, Supplemental Table S3).

**Figure 2:**
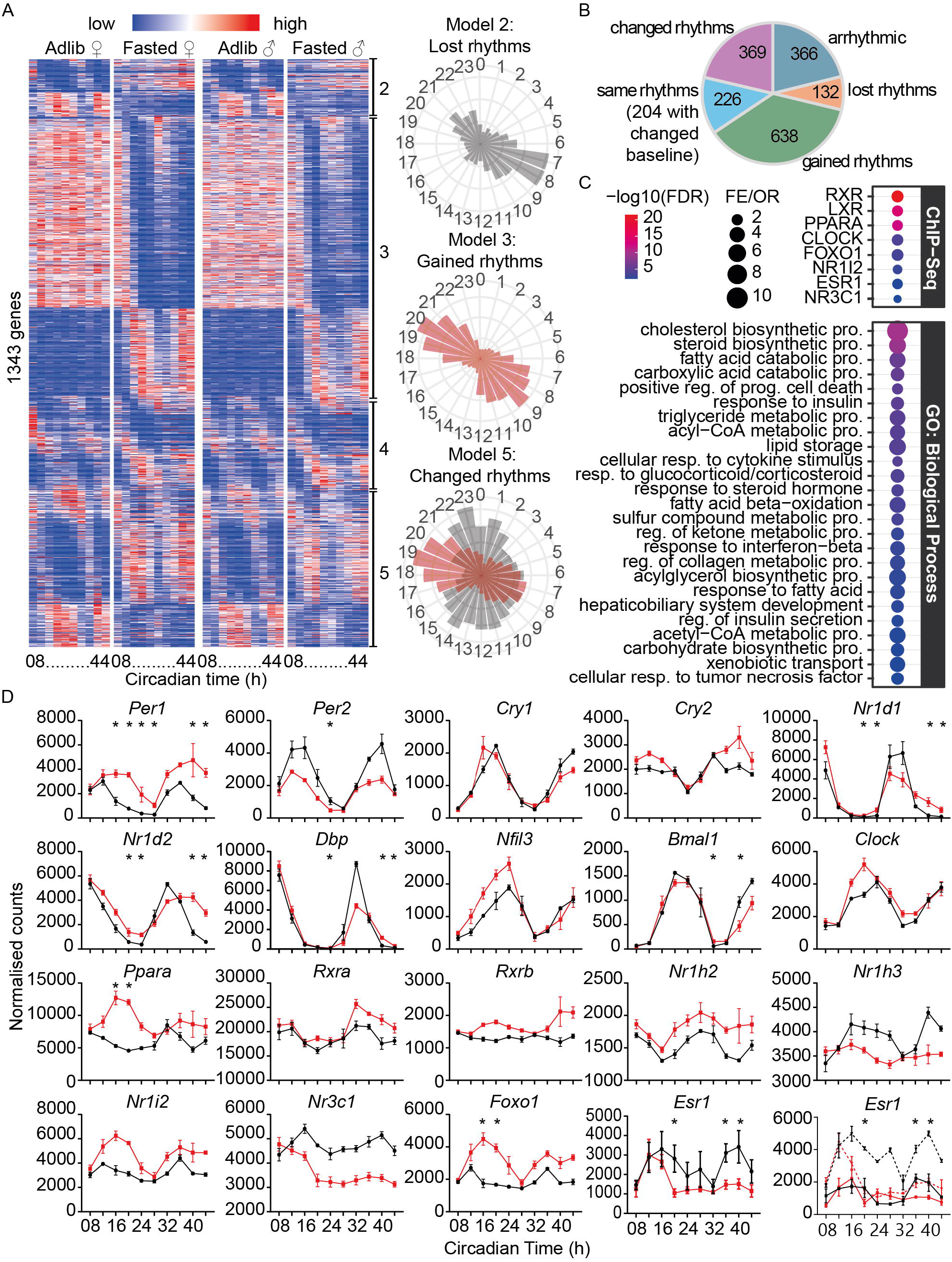
Circadian response of the hepatic transcriptome to fasting. **A**, Heatmap of high-confidence genes passing all significance thresholds (DESeq2: q < 0.05, |log2FC| > 1 in ≥2 time points; dryR: BICW > 0.8), clustered by phase in *ad libitum* livers (models 2 and 5) or fasted livers (model 3). Colours represent the mean of N = 2 males or 2 females; DESeq2 and dryR were performed with N = 4 mice (2♂ + 2♀). Polar plots show phase and amplitude distributions of genes assigned to models 2, 3 or 5; grey bars, *ad libitum*; red bars, fasted. Arrhythmic genes (model 1) and model 4 polar plot are not shown because they are not informative as their phase and amplitude are essentially unchanged. **B**, Number of genes assigned to each dryR model (BICW > 0.8) after DESeq2 filtering. Of model 4 genes, 204/226 had a different baseline. **C**, Top Enrichr (top) and PANTHER GO (bottom) results for genes in models 2, 3 and 5. **D**, Expression profiles of selected clock genes and upstream transcription factors. Data show mean ± SEM of N = 4 mice, except *Esr1* (bottom right), shown as mean ± SD for N = 2 males (—) and N = 2 females (− −); red, fasting; black, *ad libitum*. * = q < 0.05 and |log2FC| > 1 in pairwise DESeq2 with diet as primary factor and sex as secondary factor. See also Supplemental Table S1 and S3.

Strikingly, contrary to prior reports^8,9^, the absence of feeding–fasting cycles increased rather than decreased the number of rhythmic genes, rising from 727 in *ad libitum* to 1,233 in fasted livers (Figure 2B, Supplemental Table S3). This finding indicates that fasting does not dismantle the hepatic clock but instead engages a new set of clock-controlled genes, effectively adding a gear to the circadian oscillator. Extending the analysis to all genes irrespective of DESeq2 filtering (BICW > 0.8 only) further reinforced this conclusion, with rhythmic gene numbers rising from 4,071 to 5,350 under fasting (Supplemental Table S3).

GO and upstream regulator analyses of the 1,139 genes assigned to models 2, 3 and 5 (Figure 2C, Supplemental Table S3)—model 4, which shows no change in rhythmic architecture, was excluded—recapitulated the metabolic themes identified by maSigPro. These analyses additionally revealed the functional consequences of circadian reprogramming: enrichment of “cholesterol biosynthetic process” and “fatty acid beta-oxidation” marks a switch from lipid synthesis to catabolism; “xenobiotic transport” indicates rhythmic adjustment of detoxification capacity; and “response to insulin/steroid hormones” suggests that the reprogramming is driven by fasting-induced shifts in systemic hormonal signalling. ChIP-Seq upstream analysis confirmed control by metabolic sensors (PPARA, RXR, FOXO1, PXR) together with NR3C1 (glucocorticoid receptor), providing a direct link to the “response to glucocorticoid” ontology.

Among the enriched upstream regulators, CLOCK—the core circadian activator that, with its heterodimer BMAL1, constitutes the positive limb of the transcriptional clock^29^—was identified (Figure 2C, Supplemental Table S3). Its presence, together with the broad gain of rhythmicity, is consistent with a fasting-driven re-gearing of the oscillator itself. Examination of individual clock gene expression profiles revealed a diverse array of responses (Figure 2D, Supplemental Table S3). *Per1*, critical for clock entrainment, showed the most rapid and pronounced induction after food withdrawal and was classified in model 5 by dryR. *Rev-erbα* (*Nr1d1*), *Rev-erbβ* (*Nr1d2*) and *Dbp* also fell into model 5, primarily exhibiting reduced rhythmic amplitude; *Rev-erbβ* additionally showed a significant phase shift. By contrast, *Bmal1, Clock, Cry1, Cry2 and Nfil3* were allocated to model 4 by dryR, despite *Per2* and *Bmal1* meeting DESeq2 thresholds at specific time points. The expression patterns of *Bmal1, Dbp, Cry1*, and *Rev-erbα* remained largely unaffected during the initial 20 hours of food deprivation, consistent with a clock that shifts progressively rather than reacting acutely to nutrient withdrawal (Figure 2D, Supplemental Table S3).

Of the upstream transcription factors identified by Enrichr, *Ppara, Rxrb, Nr1h3, Nr1i2, Nr3c1* and *Foxo1* were classified by dryR in models 2, 3 or 5 (BICW > 0.8), though only *Ppara* and *Foxo1* also met DESeq2 thresholds. *Rxra, Nr1h2* and *Esr1* retained the same rhythmic profiles in both conditions, but only *Esr1* met DESeq2 criteria (Figure 2D, Supplemental Table S3). The markedly higher *Esr1* variability under *ad libitum* conditions reflected elevated expression in females and a pronounced fasting-induced decrease exclusive to females (Figure 2D, Supplemental Table S3).

This sex-specific *Esr1* pattern prompted a sex-stratified dryR analysis (Figure S2, Supplemental Table S3). Genes with altered rhythmicity in at least one sex were identified and visualised as heatmaps ordered by female dryR results to facilitate comparison. The number of genes in models 2, 3 and 5 was 10% higher in females (1,130) than in males (1,012), with poor overlap between sexes and relative to the pooled analysis (Figure S2C–D, Supplemental Table S3), indicating that a substantial portion of the circadian fasting response is sexually dimorphic. Polar plots revealed that genes gaining rhythms showed similar phase distributions between sexes, whereas genes losing or changing rhythms showed appreciable phase divergence (Figure S2E). Circadian amplitude was significantly higher in females under both dietary conditions and was reduced by fasting in males but not in females (Figure S2F, Supplemental Table S3).

Sex-stratified GO analyses identified a shared metabolic core encompassing fatty acid, cholesterol and steroid metabolism and responses to insulin, cytokines and glucocorticoids, consistent with the pooled analysis (Figure S2G, Supplemental Table S3). Sex-specific ontologies were less significant overall but biologically informative: “circadian rhythm” and “response to interferon-beta” were female-specific; “response to corticosteroid” and “response to TNF” were male-specific. ChIP-Seq mining revealed minimal sex differences in putative upstream transcription factors, with the exception of NR3C1, which was significant only in males (Figure S2H, Supplemental Table S3), suggesting that fasting-induced stress is transduced through distinct transcriptional intermediaries in each sex.

Collectively, these findings demonstrate that the hepatic fasting response is inextricably linked to—and actively reshapes—the circadian transcriptome. Rather than disrupting clock-driven gene expression, nutrient deprivation engages a new cohort of rhythmic genes while selectively dampening others, revealing a robust endogenous control over the temporal organisation of metabolic priorities. The female-specific enrichment of circadian-related ontologies suggests tighter coupling between the molecular oscillator and the nutritional response in females, whereas the male response appears more strongly influenced by extrinsic glucocorticoid signalling.

### Alternative transcriptome response to fasting

Beyond changes in transcript abundance, the temporal coordination of the transcriptome may be further diversified by post-transcriptional mechanisms. Differential exon usage—a key source of proteome diversity in eukaryotes^30^— represents a potential secondary regulatory layer that is independent of bulk mRNA levels. To characterise this layer, we extended our analysis to exon-level quantification using DEXSeq^31^.

A progressive effect of fasting on exon usage was evident from the number of exons meeting significance thresholds (q < 0.05, |log2FC| > 0.58) at each time point: 76 at CT08, 123 at CT12, and between 1,010 and 1,759 at subsequent time points (Figure 3A–B, Supplemental Table S4). The low and inconsistent numbers at CT08 and CT12—before or immediately after fasting onset—likely reflect stochastic variation. From CT16 onwards, between ~600 and 1,200 exons were significant at a single time point only, suggesting a combination of biologically meaningful time-dependent variation and nutrient-deprivation-induced stochastic splicing events.

**Figure 3:**
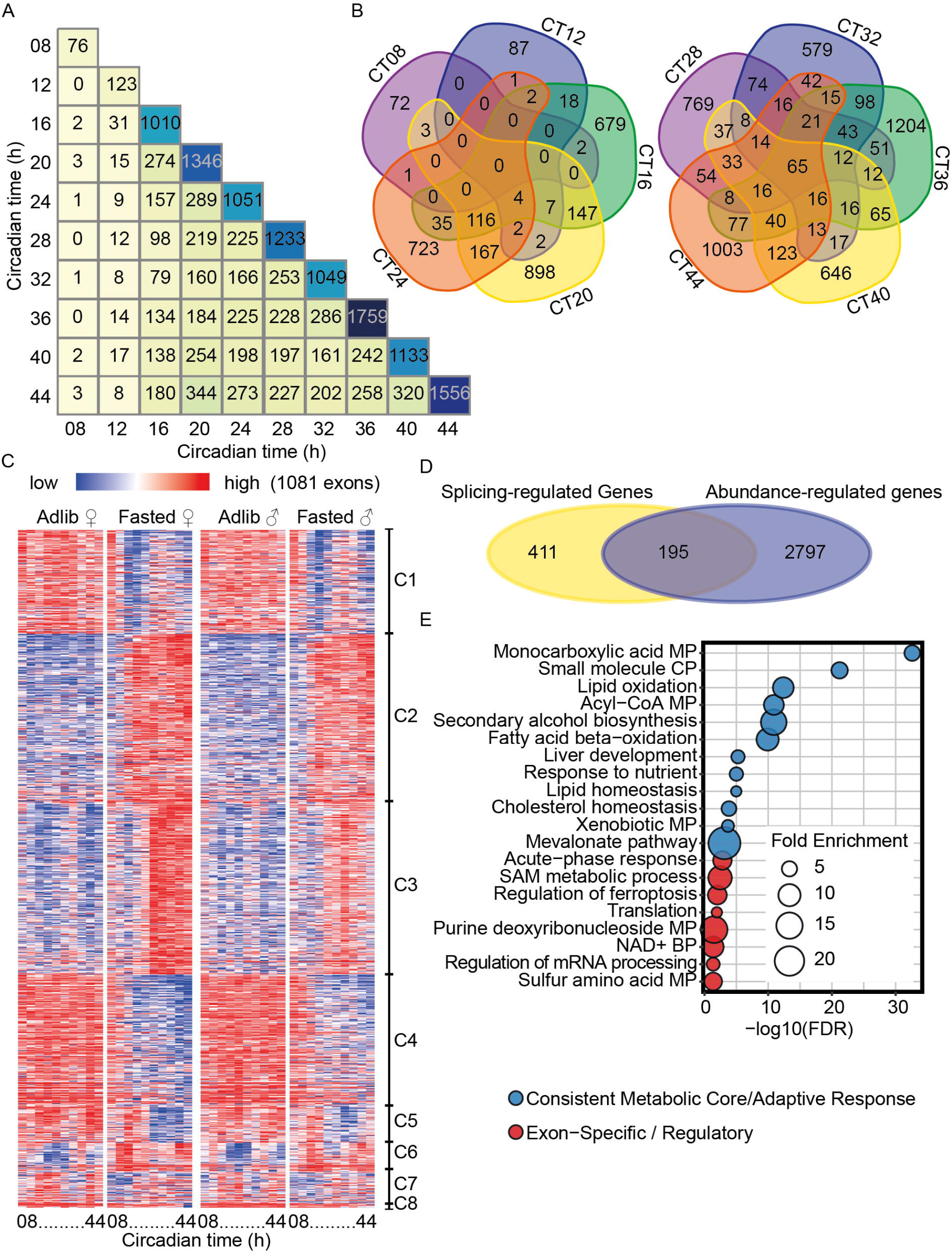
Phased alternative splicing transitions and kinetic landscape of the hepatic fasting response. **A** and **B**, Number and overlap of exons significant (q < 0.05, |log2FC| > 0.58) by DEXSeq at each time point. **C**, Heatmap of high-confidence exons passing all selection thresholds (DEXSeq: adjusted *p* < 0.05, |log2FC| > 0.58 in ≥2 time points; maSigPro: *p* < 0.05), clustered by usage kinetics. Colours represent the mean of N = 2 males or 2 females; DEXSeq and maSigPro were performed with N = 4 mice (2♂ + 2♀). **D**, Venn diagram of the overlap between genes regulated at the expression level and at the exon usage level. **E**, Selected PANTHER GO results for the 606 genes in C. MP, metabolic process; BP, biosynthetic process; CP, catabolic process. See also Supplemental Table S5.

Integrating DEXSeq with maSigPro trend analysis identified 1,081 high-confidence exons from 606 genes, organised into eight clusters (Figure 3C, Supplemental Table S5). The four largest clusters (1–4) capture the dominant splicing transitions: Cluster 1, characterised by decreased exon usage from CT16 with a superimposed rhythmic component; Clusters 2 and 3, both showing increased usage with distinct kinetics; and Cluster 4, showing a more linear decrease. Crucially, only ~one third of the 606 genes (195) were also regulated at the transcript abundance level (Figure 3D), demonstrating that differential exon usage constitutes an independent layer of fasting-induced regulation.

GO analysis of the 606 genes revealed that, while a consistent metabolic core—fatty acid oxidation, ketone body and xenobiotic metabolism—is regulated at both the transcript and exon levels, several highly specific biological processes emerged exclusively among genes regulated through exon usage (Figure 3E, Supplemental Table S5). Most notably: S-adenosylmethionine (SAM) metabolism (11.53-fold enrichment), purine deoxyribonucleotide metabolism (15.81×), NAD^+^ biosynthesis (8.68×), ferroptosis (7.14×), translation (2.22×) and mRNA processing (2.89×). This pattern indicates that while the liver upregulates global energy mobilisation at the transcriptional level, it deploys alternative splicing to fine-tune cofactor biosynthesis, cellular quality control and the mRNA processing machinery itself.

A sexual dimorphism was evident in exon usage, with changes more pronounced in females (Figure 3C, Supplemental Table S5). Sex-stratified maSigPro analysis identified 702 high-confidence exons in females versus 483 in males, with 361 shared (Figure S3A–C, Supplemental Table 5). Only 21 female-specific and 15 male-specific exons were detected beyond those found in the pooled analysis, again indicating that the dimorphism is primarily quantitative. These 36 exons spanned 31 genes, including female-specific differential usage of *Esr1*, the adipose triglyceride lipase *Pnpla2* and several cytochrome P450 genes (*Cyp3a25, Cyp2c69, Cyp2a22*), and male-specific usage of *Ptch1* and *Selenbp2* (involved in hormone signalling and detoxification, respectively; Figure S3D, Supplemental Table S5).

A rhythmic component in exon usage was apparent in Clusters 1, 3 and 6 (Figure 3C), prompting us to formally test whether exon usage exhibits circadian rhythmicity—a question not previously addressed—and whether fasting alters these rhythms. Applying dryR to exon-level data, we identified 382 high-confidence exons from 227 genes with fasting-dependent changes in circadian rhythmicity (Figure 4A, Supplemental Table S6): 164 gained rhythmicity, 62 lost rhythmicity, 55 showed altered rhythms, and 101 with unchanged rhythmic architecture but a shifted mean usage level across the time course (model 4; Figure 4A). Exons gaining rhythms (model 3) mostly peaked around CT08 or CT20 in fasted livers, reflecting antiphasic oscillations that emerge during fasting (Figure 4A, Supplemental Table S6). These observations establish that exon usage, like transcription, is subject to circadian clock control, and that fasting drives a gain of circadian rhythmicity at the post-transcriptional level in parallel with its effects at the transcript level.

**Figure 4:**
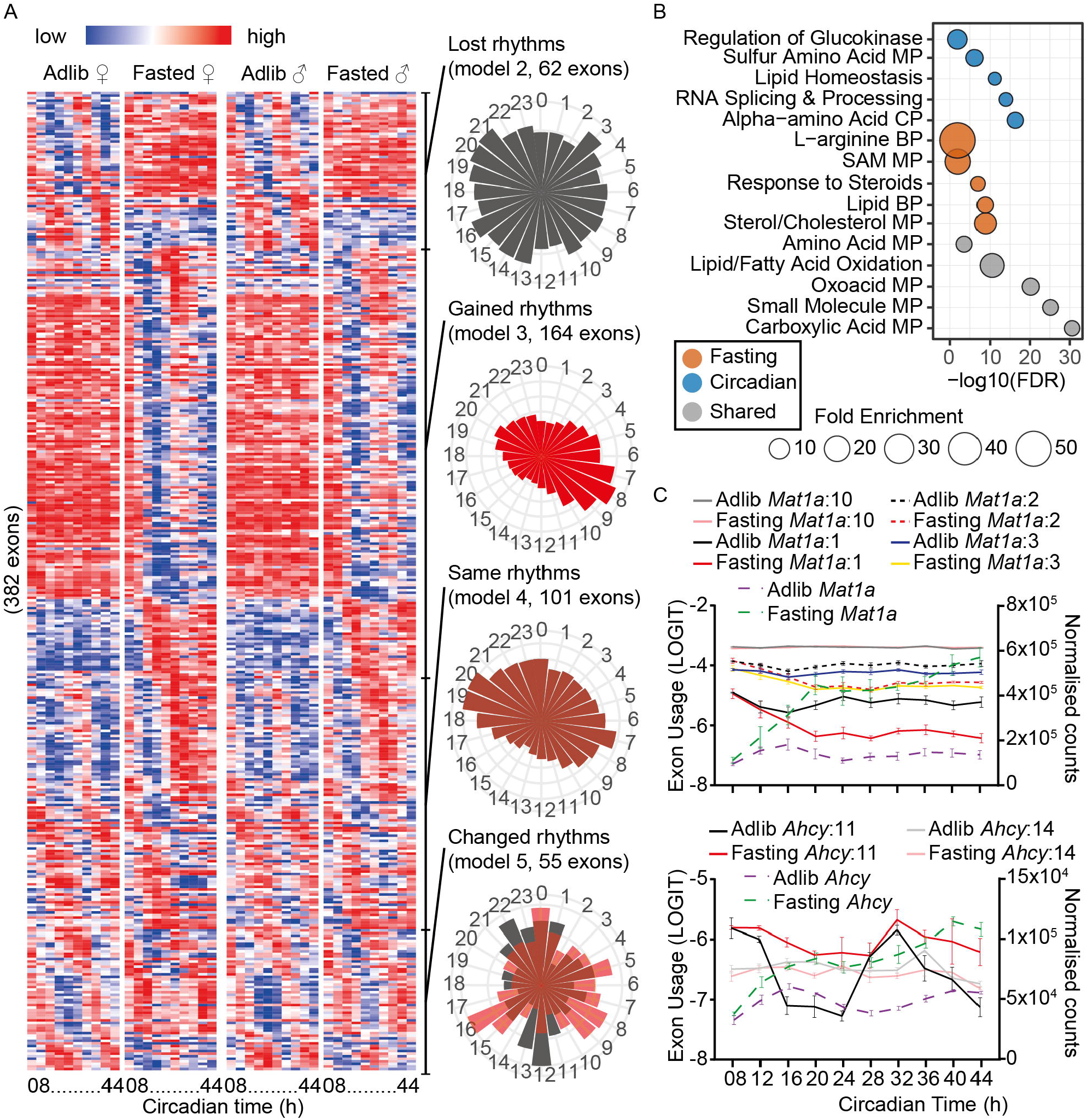
Circadian response of the hepatic alternative transcriptome to fasting. **A**, Heatmap of high-confidence exons passing all selection thresholds (DEXSeq: adjusted *p* < 0.05, |log2FC| > 0.58 in ≥2 time points; dryR: BICW > 0.8), clustered by phase in *ad libitum* livers (models 2, 4 and 5) or fasted livers (model 3). Colours represent the mean of N = 2 males or 2 females; DEXSeq and dryR performed with N = 4 mice (2♂ + 2♀). Polar plots on the right show phase and amplitude distributions for models 2, 3, 4 and 5; grey bars, *ad libitum*; red bars, fasted. Here we showed the polar plot for model 4 exons to appreciate their non-random phase diversity, as this has not been reported before. Plots with arrhythmic exons (model 1) are not shown. **B**, PANTHER GO results from two parallel analyses: “Fasting” group (227 genes from 382 exons in A) and “Circadian” group (1,990 genes with rhythmically used exons in *ad libitum* livers, BICW > 0.8, unfiltered by DEXSeq). Shared ontologies are indicated. MP, metabolic process; BP, biosynthetic process; CP, catabolic process; SAM, S-adenosylmethionine. **C**, Expression and exon usage profiles of *Mat1a* and *Ahcy* during fasting. Exon numbers follow genomic coordinates (left-to-right); gene name alone indicates transcript abundance (right axis). Invariable exons shown for reference: exon 10 (*Mat1a*), exon 14 (*Ahcy*). See also Supplemental Table S6.

Two parallel GO analyses were performed: one with the 227 genes showing fasting-dependent changes in circadian exon usage (“Fasting” group) and one with the 1,990 genes carrying rhythmically used exons under *ad libitum* conditions (“Circadian” group; Figure 4B, Supplemental Table S6). The Circadian group highlighted constitutive roles of alternative splicing in hepatic clock function across metabolic pathways from amino acid to lipid metabolism, and in the regulation of mRNA splicing itself. Notably, the SAM metabolic process emerged as highly enriched in the Fasting group, consistent with the previous GO enrichment based on maSigPro’s results and further highlighting the connection between methionine/SAM metabolism, biological methylation and circadian clock function ^32–36^ (Figure 4B, Supplemental Table S6).

Critically, among genes annotated to the SAM metabolic process ontology, *Mat1a* and *Ahcy*, the two core enzymes of the methionine/methyl cycle, were both flagged as regulated by splicing by maSigPro and/or dryR. MAT1A synthesises SAM from ATP and methionine in the liver; the resulting SAH must be rapidly hydrolysed to homocysteine by AHCY to prevent competitive inhibition of downstream methyltransferases. *Mat1a* and *Ahcy* were also identified among the 2,995 fasting-responsive genes in Cluster 3 (Figure 1A), and their total transcript levels rose progressively and substantially with fasting duration (Figure 4C). Fasting simultaneously triggered distinct changes in the temporal profile of specific exons: the usage of three *Mat1a* exons—together defining alternative transcriptional start sites and distinct 5’-UTRs (Figure S4)—declined after 10 hours of fasting and showed a subsequent weak recovery, classified as a gain of rhythmicity (model 3); one *Ahcy* exon (exon 11, a retained intron) was significantly upregulated in fasted liver and showed attenuated circadian amplitude, classified in model 5 (Figure 4C). These splicing events are reminiscent of the METTL16-dependent regulation of *Mat2a*—the liver-excluded paralogue of *Mat1a*—where intron retention is coupled to cellular SAM availability^37,38^. Whether analogous m^6^A-dependent mechanisms operate on *Mat1a* and *Ahcy* in hepatocytes remains to be determined, but these genes exemplify a class of metabolic regulators where fasting re-gears the circadian programme of isoform selection independently of bulk transcriptional output.

Sex-stratified dryR analysis of circadian exon usage revealed low overlap between males and females in both exons and genes (Figure S5A, Supplemental Table S6): 224 exons from 145 genes in females; 301 exons from 177 genes in males; 95 exons (74 genes) shared. Phase distributions of rhythmic exon usage were sexually dimorphic even under *ad libitum* conditions (Figure S4D), whereas amplitude differences were modest and not significant (Figure S5B–C). GO analyses with male and female datasets revealed a striking complementarity: sex-unique transcriptional responses were twice as prevalent in females (Figure S2), whereas sex-unique alternative splicing events were twice as prevalent in males (Figure S4E, Supplemental Table S6). Furthermore, while transcriptional dimorphism spanned broad hormonal and signalling pathways, dimorphism in exon usage was concentrated in core metabolic machinery.

In summary, the hepatic response to fasting operates through a hierarchy of temporal regulatory layers. Total transcript abundance drives a global shift toward fat-burning and ketone production, while alternative exon usage provides an independent, nuanced layer of control specifically targeting cofactor biosynthesis, cellular quality control and mRNA processing. Fasting does not merely perturb the hepatic transcriptome: it actively re-gears it, inducing new circadian rhythms in both gene expression and splicing. The fundamental metabolic blueprint is conserved between sexes, with sexual dimorphism arising primarily from the magnitude and rhythmic robustness of the response, and — at the level of circadian splicing — the regulatory strategies employed.

## DISCUSSION

This study provides a high-resolution, multi-layered atlas of the hepatic fasting response, demonstrating that the transition from nutritional abundance to deprivation is not a passive depletion of resources but an actively orchestrated re-gearing of the liver’s molecular machinery. By integrating temporal transcriptomics with exon-level analysis across both sexes over a continuous 36-hour window, we show that this re-gearing is governed by a regulatory hierarchy in which transcription and alternative splicing cooperate—and are both subject to circadian control—to drive metabolic adaptation.

A central finding is that alternative splicing operates as a fundamental regulatory layer for hepatic metabolic control, acting independently of total mRNA abundance. Our analysis identifies specific processes—most notably the SAM/methionine cycle—where the fasting response is primarily re-geared through changes in the temporal profile of exon usage rather than overall transcript levels. The isoform shifts in *Mat1a* and *Ahcy* are particularly instructive: while total transcript levels for both enzymes rise progressively with fasting—consistent with increased demand for methyl-donor metabolism—concurrent fasting-induced changes in the circadian rhythmicity of specific exons reveal an additional, independent layer of regulation operating at the level of isoform selection. Studies relying solely on transcript abundance may therefore overlook a critical post-transcriptional rheostat fine-tuning the activity of key metabolic bottlenecks. This principle extends to the circadian field: Koike *et al*. (2012)^39^ demonstrated that distinguishing nascent from mature transcripts within the same RNA-seq dataset reveals that transcriptional and post-transcriptional regulation make independent contributions to rhythmic gene expression in the liver — underscoring that the regulatory complexity uncovered here through exon-level analysis was always present but invisible to bulk transcript measures.

The enrichment of ferroptosis-related genes among those showing fasting-induced changes in exon usage is particularly noteworthy: this form of iron-dependent, lipid peroxide-driven cell death was largely absent from the transcript abundance analysis, suggesting that hepatic defences against lipid peroxidation during nutrient deprivation are regulated primarily at the isoform level. More broadly, the finding that fasting induces new circadian rhythmicity in exon usage—including in RNA splicing and mRNA metabolic processes themselves—indicates that the hepatic clock does not merely oscillate transcript levels but actively cycles isoform selection to maintain metabolic homeostasis across the 24-hour day, independently of immediate nutritional cues.

By including both sexes, we have uncovered a hierarchy of sexual dimorphism across regulatory layers. The “core” circadian splicing machinery is relatively conserved, while the more plastic responses to fasting—gene expression and non-circadian splicing—diverge between sexes. Females exhibit more robust transcriptional response, whereas males deploy a broader repertoire of alternative splicing events, suggesting that the two sexes achieve broadly similar metabolic ends through distinct regulatory toolkits. The discovery that circadian splicing dimorphism resides primarily in phase distribution (timing) rather than amplitude (magnitude) suggests that male and female livers follow distinct temporal schedules for core metabolic processes—a distinction that may underlie known sex differences in susceptibility to metabolic disorders upon circadian disruption^40–44^.

The extent of this dimorphism depends critically on the analytical approach. maSigPro—whether applied to pooled or sex-separated data—identified dimorphism as primarily quantitative and marginal in scope. By contrast, dryR uncovered a far richer landscape, revealing substantial sex differences in both amplitude and phase of rhythmic gene expression that were evident even under *ad libitum* conditions, demonstrating that the basal hepatic circadian transcriptome—including alternative splicing—is constitutively and inherently dimorphic^41^. This discrepancy highlights a fundamental limitation of standard longitudinal trend modelling: maSigPro captures the magnitude of a transcriptional response but lacks the sensitivity to detect phase shifts that define sex-specific chronobiology. Metabolic studies often fail to account for circadian responses, and circadian studies often fail to account for non-circadian responses; the present work integrates both.

The identification of CLOCK as a significant upstream regulator of the late-stage progressive response (Cluster 5, Supplemental Table S2), together with its enrichment among genes gaining rhythmicity during fasting (Figure 2C), suggests a temporal hierarchy in the re-gearing of the circadian oscillator: the liver first mounts an acute gluconeogenic response driven by GR, then more protractedly activates the ketogenic programme as GR induces PPARA expression — a cascade directly consistent with the enhancer dynamics previously described^45^ — before ultimately restructuring the clock machinery itself. This provides a mechanistic explanation for apparent discrepancies with prior work reporting fewer rhythmic genes after prolonged fasting^8,9^: those studies likely sampled the liver after the initial re-gearing had stabilised into a new hypometabolic steady state. Our 36-hour time course captures the dynamic transition during which the oscillator is actively being re-geared, before that stabilisation occurs. The apparent “loss” of rhythmicity observed elsewhere may therefore reflect late-stage clock consolidation rather than clock disruption.

The raw high-depth sequencing dataset generated here—encompassing both sexes across a 36-hour time course at 4-hour resolution—constitutes a comprehensive resource for the metabolic research community, enabling meta-analysis of sex-specific gene–environment interactions and exploration of isoform-specific metabolic regulation that are inaccessible in smaller or single-sex studies.

### Limitations and Future Directions

The principal limitation of this study is its exclusive reliance on RNA-level data. The downstream consequences of the transcriptional and splicing shifts identified here— with respect to protein isoform abundance, enzymatic activity and metabolic flux— remain to be validated. Future work should prioritise proteomic and metabolomic profiling across a comparable time course, with particular attention to the pathways identified as targets of isoform-level regulation, especially the methionine cycle and ferroptosis-related networks. The sex-stratified maSigPro and dryR analyses rest on N = 2 mice per sex; however, the inclusion of sex as a covariate in the DESeq2 and DEXSeq pairwise models means that the primary expression and exon usage results are not subject to this limitation. Furthermore, the high temporal density of the dataset—ten time points spanning 36 hours—provides degrees of freedom that partially compensate for the modest per-sex replicate number in the trend and rhythmicity analyses, though confirmation in larger sex-balanced cohorts remains desirable. We mitigated the inherent risk of detecting statistically significant trends of negligible biological effect by requiring all reported genes/exons to additionally meet stringent fold-change thresholds in independent pairwise comparisons. Finally, while long-read sequencing would offer advantages in directly resolving full-length isoforms, the depth and temporal density of our short-read dataset (80 samples across 36 hours) provides the replicate power to validate exon-level changes across consecutive time points, reducing the risk that stochastic splicing events are reported as biologically meaningful.

In conclusion, the hepatic response to fasting is a multi-dimensional process in which transcription and alternative splicing cooperate—and are both subject to circadian re-gearing—to drive coordinated metabolic adaptation to nutrient deprivation. Sexual dimorphism in this response is real but primarily quantitative: a difference in the magnitude and temporal precision of a shared programme rather than a fundamental divergence in regulatory logic. Understanding how the liver synchronises its molecular machinery with both time of day and nutritional state, and how this integration differs between sexes, will be essential for developing sex-specific therapeutic strategies for metabolic syndrome and related nutritional pathologies. The multi-layered, open-access atlas presented here provides a foundation for that endeavour.

## METHODS

### Animals and liver sampling

Animal experiments were licensed under the Animals (Scientific Procedures) Act of 1986 (UK) and were approved by the animal welfare committees at the University of Manchester. Nine-to twelve-week-old C57BL/6J male and female mice were purchased from Charles River (UK) and acclimatized to the local animal unit for 1 week before the experiment, with daily inspection and handling. Mice were housed by groups of 4 under a 12h:12h light/dark cycle at ~250 lux during the light phase and 0 lux during the dark phase; time of lights on 8:00, lights off 20:00. Mice had free access to nesting material and a cardboard tunnel. Ambient temperature was kept at 22□±□3□°C, the humidity was ~52□±□4%, with food (Special Diets Services, BK001) and water available *ad libitum*. At the beginning of the first day in constant darkness, all mice were transferred to wire-bottomed cages to limit coprophagy. At 16:00 on the first day in constant darkness (CT08), the liver samples from the first two groups of 4 mice were sampled and immediately snap frozen in liquid nitrogen. At 18:00 on the first day in constant darkness (CT10), food was removed from a random half of the remaining 18 cages. At 20:00 on the first day in constant darkness (CT12) and every four hours thereafter until CT44, liver samples from one group of *ad libitum*-fed and one group of fasted mice were collected and immediately snap frozen in liquid nitrogen. All liver samples were collected from the central, thickest part of the left lobe and weighed ~50mg. Samples were kept at −80°C until use.

### RNA extraction

One by one, each liver sample was promptly transferred to a 2ml microcentrifuge tube (Starlab I1420-2702) kept at 4°C and containing 1 ml TRIzol (ThermoFisher 15596018) and homogenized with a Polytron (Kinematica PT1200E) for 1-2 minutes until complete lysis, followed by a 5-minute incubation at 4°C. The homogenizer head was thoroughly washed and dried between each sample. Total RNA was then extracted following standard TRIzol protocol.

### Bulk RNASeq and gene expression analyses

Total RNA was submitted to the Genomic Technologies Core Facility (GTCF) of the University of Manchester for bulk RNA sequencing as previously described^33^. Reads were aligned to the mouse genome Gencode Release M38 (GRCm39) using HISAT2 with Gencode annotations (gencode.vM38.annotation.gtf), and genes were counted using featurecounts. To identify high-confidence regulated genes, a two-pronged approach was used. Pairwise DESeq2^15^ analyses were first performed for each timepoint, with outlier filtering using Cook’s distance and independent filtering using the mean of normalized counts but without pre-filtering for low count genes, with diet (*ad libitum* vs fasting) as primary factor and sex (male vs females) as secondary factor. Raw counts, normalized counts and DESeq2 results are available in Supplemental Table S1.

Next, to identify gene affected by fasting in a time-dependent manner, we used Method for Analysis of SIGnificant PROfiles (maSigPro)^13,14^, an R package that uses a two-step regression strategy to identify differentially expressed genes and then uses stepwise regression to find clusters of genes with similar temporal expression profiles. We limited this analysis to genes having at least 10 raw counts (counted by featurecounts) in at least 20 of the 80 samples (25%), but counts normalized by the median of ratios were used as input for maSigPro. The resulting input dataframe contained 18,023 genes out of a total of 78,275 genes in the annotations that included known genes as well as gene models. The output results from maSigPro include several p values corresponding to the overall significance of the model (“p value”), regression coefficients significantly different from 0 (p.value_time…) that may be significantly different between groups (p.value_timexfasting). While maSigPro reports the nominal p values in the output, the significant genes or exons were filtered using a threshold q value < 0.05 of (Benjamini-Hochberg)^13,14^.

To identify genes whose circadian expression is affected by fasting, we analysed the same subset of 18023 genes using dryR^28^ (but using raw counts instead of normalised counts), a R package that uses a Bayesian model selection framework for analysing differential rhythmicity in RNAseq data, allowing for the comparison of rhythmic patterns (amplitude and phase) and mean expression levels across two conditions. DryR groups genes across 5 models: arrhythmic in both conditions (model 1), losing rhythms under fasting (2), gaining rhythms in fasting (3), identical rhythms in both conditions (4) and with changed rhythms (5). To identify high-confidence genes showing different circadian rhythms of expression with high confidence level of the selected dryR model, we limited our analysis to genes with a Bayesian Information Criterion Weight (BICW) of over 0.8.

To further ensure that these statistically significant shifts in time-dependent profiles (maSigPro) and circadian rhythmicity (dryR) represented functionally meaningful biological signals, we cross-referenced these results with our pairwise DESeq2 analysis. Only those genes with significant shifts in trends with maSigPro, or categorised by dryR as having altered rhythms (Models 2, 3, or 5), and that also met a stringent biological threshold—defined as an adjusted *p* value < 0.05 and a |log2 fold-change| > 1 by DESeq2 in at least two time points—were considered high-confidence candidates for downstream functional enrichment and included in the main figures. This dual-filtering approach effectively prioritises genes where the nutritional challenge of fasting triggers a robust reorganisation of the temporal transcriptome rather than subtle or stochastic fluctuations in expression levels.

### Exon usage analyses

First, we used DEXSeq^31^ count function to obtain read counts for all exons in each samples, using gencode.vM38 annotation file previously prepared by DEXSeq for counting. As with gene expression, to identify high-confidence regulated exons, pairwise DEXSeq analyses were first performed for each timepoint, with diet (*ad libitum* vs fasting) as primary factor and sex (male vs females) as secondary factor. Raw unfiltered exon counts and DEXSeq results are available in Supplemental Table S7.

Next, exon count data was pre-processed to allow analyses with dryR and maSigPro. Exons with low counts were first removed, keeping exons with at least 10 counts in at least 20 samples (1/4 of the total), before calculating exon usage for each exon for each gene (exon count/total gene count). To enable further calculations, exon usage values equal to 0 were replaced by the smallest non-zero exon usage value (a, floor value), exon usage values equal to 1 were replaced by 1-a (ceiling value), and exon usage values equal to 0.5 were replaced by 0.4999999. To satisfy the Gaussian assumptions of the linear modeling frameworks in both maSigPro and dryR, exon usage proportions were logit-transformed (LN(exon usage/1-exon usage)). This transformation ensured numerical stability and permitted the accurate estimation of both polynomial trends and harmonic parameters across the 36-hour time course. Lastly, a variance-based filter was applied where exons with >40 floor/ceiling values were discarded. This focused the differential usage analysis on exons with sufficient dynamic range to model temporal shifts (dryR) and containing enough non-constant data points to support the degrees of freedom required for a two-way polynomial fit (maSigPro). Furthermore, reducing the number of uninformative features decreased the multiple-testing burden, thereby increasing the statistical power to detect genuine time-dependent interactions. The resulting dataframe, containing 198,649 exons from an initial number of 470,140, was then used as input for maSigPro or dryR. We cross-referenced these exon usage results with our pairwise DEXSeq analysis. Only those exons with significant shifts in trends with maSigPro, or categorised by dryR as having altered rhythms (models 2, 3, or 5), and that also met a stringent biological threshold—defined as an adjusted *p* value < 0.05 and a |log2 fold-change| > 0.58 by DEXSeq in at least two time points were considered high-confidence candidates. We applied a distinct fold-change threshold from the one used with gene expression because with total transcriptomic abundance above, a 2-fold change threshold accounts for the high dynamic range and inherent variance of mRNA counts, but a 1.5-fold threshold is more appropriate for differential exon usage because DEXSeq coefficients represent changes in exon abundance relative to the total gene transcript pool, effectively normalising for fluctuations in overall gene expression. A 1.5-fold shift in relative usage indicates a substantial change in the molecular stoichiometry of the transcript population. Such shifts are sufficient to trigger functional consequences, such as the inclusion of premature termination codons or retained introns leading to nonsense-mediated decay^37,38^ or the alteration of critical protein domains that dictate subcellular localisation and signalling activity^46,47^.

### GO analysis and upstream Chip-Seq data mining

All GO analyses were performed by PANTHER^16^ on geneontology.org^48,49^ using the standard background gene list for *Mus musculus*. Chip-Seq data mining was performed on maayanlab.cloud/Enrichr/^18–20^ using the ChEA Transcription Factor Targets 2022 database^50,51^. Unless specifically mentioned, only transcription factors from mouse liver datasets are included in the main figures, but Supplemental Table files show the entire Enrichr output. In polar plots of GO results, sex bias is calculated as angle = atan2(-log10(P_Male_), −log10(P_Female_)).

## DATA AVAILABILITY

Raw data and associated processed files have been deposited to GEO, dataset GSE328655. Supplemental data files are publicly available via the University of Manchester Figshare repository at https://doi.org/10.48420/32091148.

## ACKNOWLEDGEMENTS

The authors would like to gratefully acknowledge the help of the Genomic Technologies Core Facility, the assistance given by Research IT and the use of the Computational Shared Facility at The University of Manchester.

## AUTHOR CONTRIBUTIONS

Jessica Treeby: Investigation, Formal analysis, Writing - Review & Editing. Shin-Yu Kung: Software. Benjamin Saer: Investigation, Methodology. Judith Gramunt Prillo: Software. Louise Hunter: Supervision, Formal analysis. Jean-Michel Fustin: Conceptualization, Methodology, Software, Formal analysis, Investigation, Data Curation, Visualization, Supervision, Project administration, Funding acquisition.

## FIGURE LEGENDS

**Figure S1:**
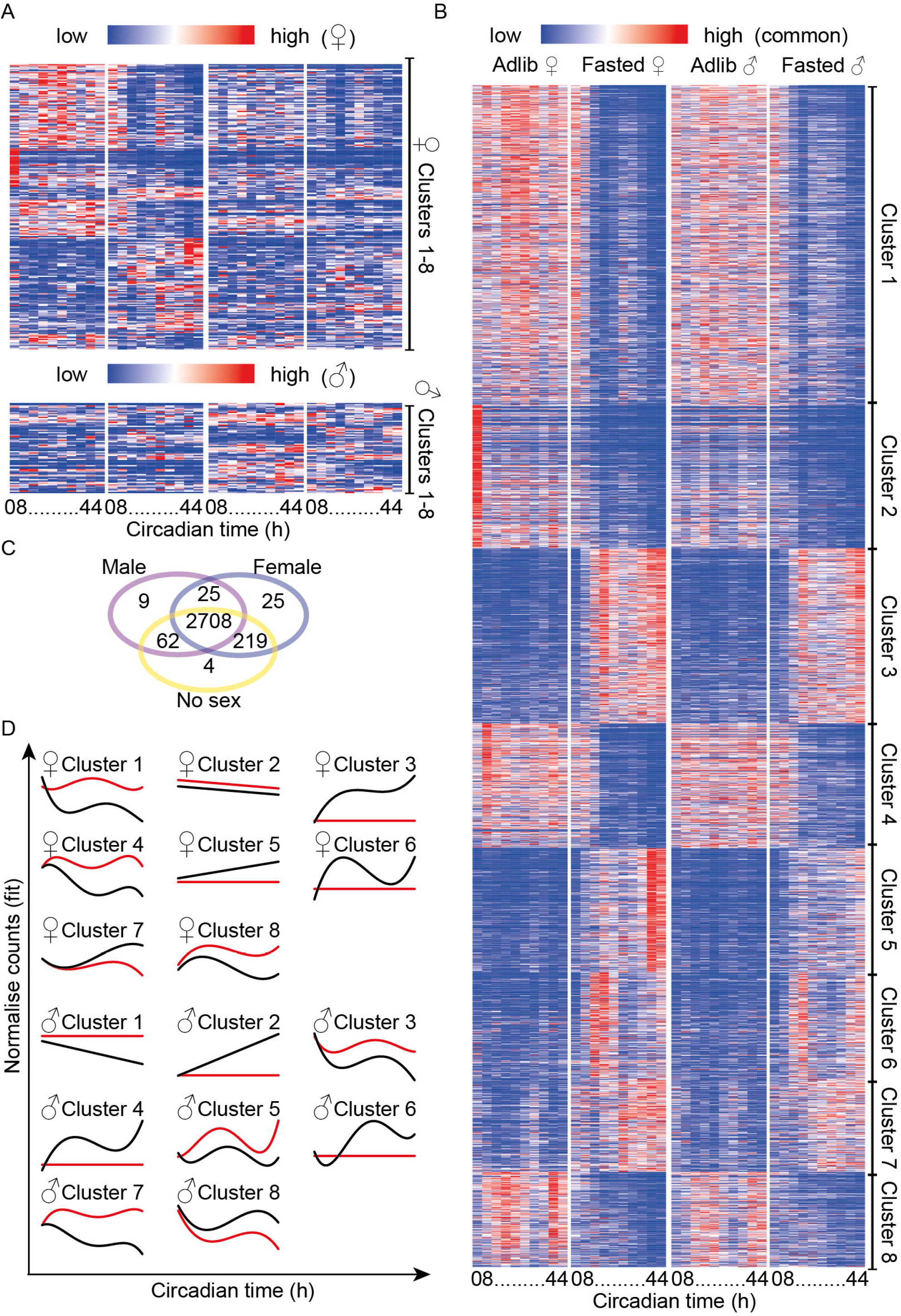
Sex differences in the phased transcriptional response to fasting. **A**, Heatmaps of high-confidence genes significant only in males or only in females (**B**, common genes). **C**, Venn diagram of genes identified by maSigPro with male-only (N = 2), female-only (N = 2), or combined (N = 4) data. **D**, maSigPro-fitted expression curves for male and female clusters (cluster numbering is automatic and independent between sexes; note similar fit for example for F1 and M3, F3 and M4, F4 and M7, F5 and M2, F7 and M8, F8 and M5). See also Supplemental Table S1 and S2.

**Figure S2:**
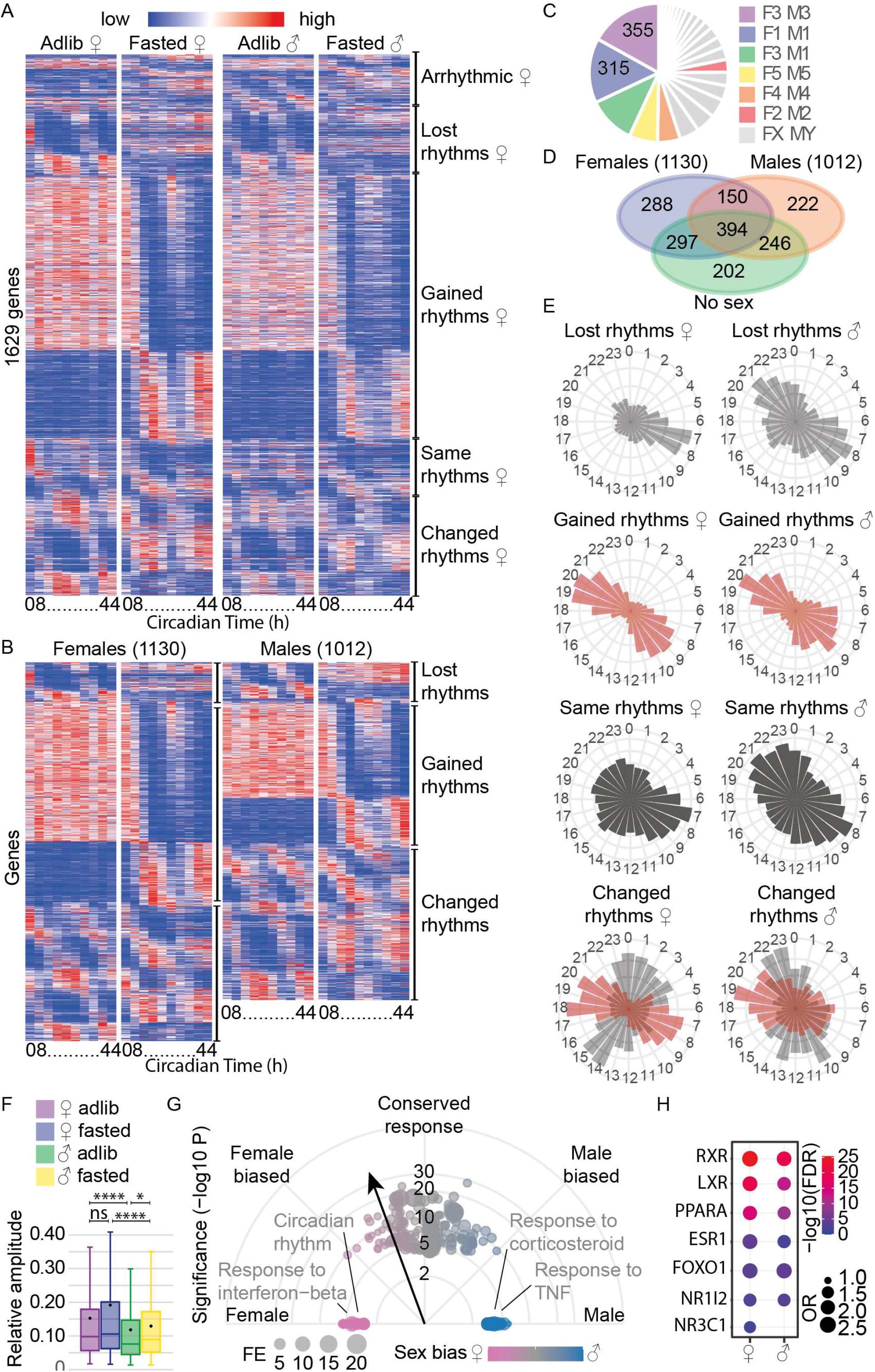
Sex-specific circadian responses of the hepatic transcriptome. **A**, Heatmap of genes with altered rhythmicity (DESeq2: q < 0.05, |log2FC| > 1 in ≥2 time points; dryR in ≥1 sex: BICW > 0.8), ordered by female dryR phase (models 2, 4 and 5: *ad libitum* phase; model 3: fasted phase; model 1: ordered similarly by male models and phase). Colours represent the mean of N = 2 males or 2 females; dryR performed independently with N = 2 per sex. **B**, As in A, but genes in models 2, 3 and 5 shown in separate male and female heatmaps ordered by their respective phases. **C**, 2D pie chart of dryR model correspondence between male (M) and female (F) data; “FX MY” denotes all remaining model combinations. **D**, Venn diagram of genes in models 2, 3 or 5 from pooled (N = 4), male-only (N = 2) or female-only (N = 2) dryR analyses. **E**, Polar plots of circadian phase differences between sexes for each model. **F**, Box-and-whisker plots of relative rhythmic amplitude (BICW > 0.8 genes) per group; dots indicate means, horizontal lines indicate medians. Kruskal–Wallis test (p < 0.0001) with multiple comparisons (*, p < 0.05; ****, p < 0.0001). **G**, Polar plot of aggregated GO results for genes in models 2, 3 and 5 in males or females. Vector length = −log10(FDR); dot size = fold enrichment; angle = sex bias (0°, female-specific; 90°, shared; 180°, male-specific). **H**, Top Enrichr upstream regulator results for male and female model 2, 3 and 5 genes. See also Supplemental Table S3.

**Figure S3:**
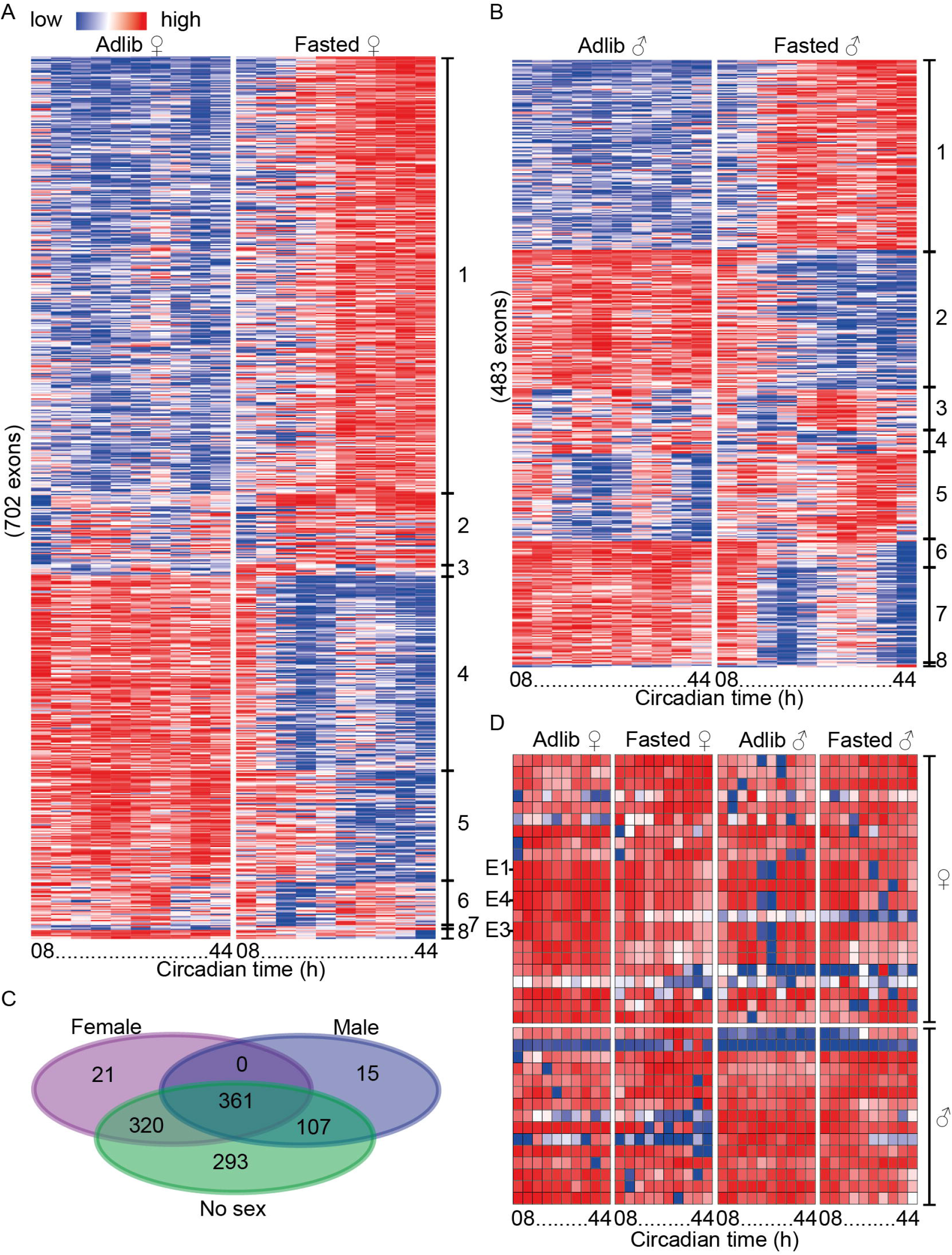
Sex differences in exon usage during fasting. Heatmaps of high-confidence exons with significant trends between *ad libitum* and fasted livers in females (**A**) or males (**B**) by maSigPro (DEXSeq: q < 0.05, |log2FC| > 0.58 in ≥2 time points). Colours represent the mean of N = 2 males or 2 females. **C**, Venn diagram of differentially used exons identified by maSigPro with female-only (N = 2), male-only (N = 2) or combined (N = 4) data. **D**, Heatmap of the 21 female-specific and 15 male-specific exons from C; exons 1, 3 and 4 of *Esr1* (significant only in females) are labelled for illustration. See also Supplemental Table S5.

**Figure S4:**
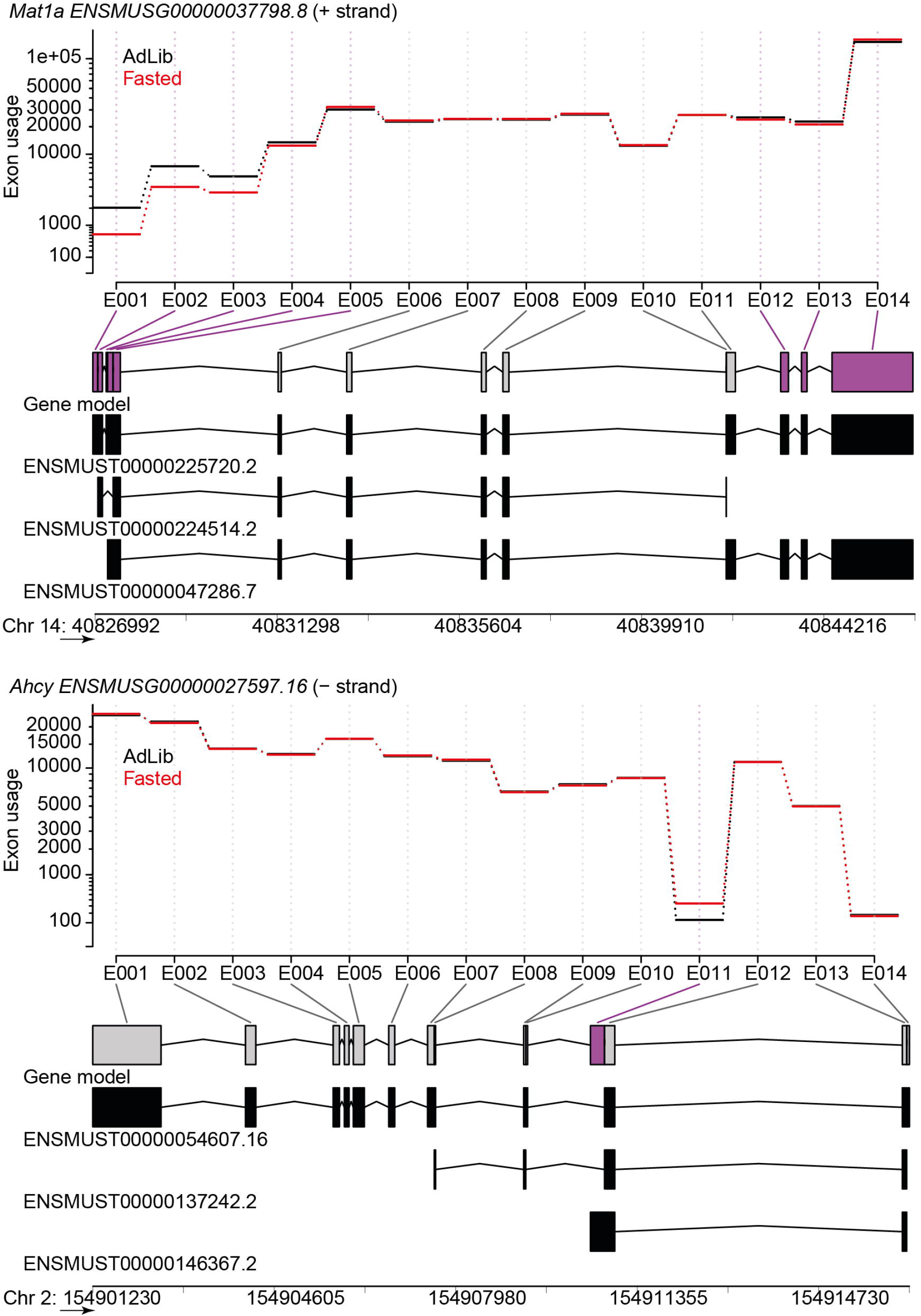
Fasting-induced alternative splicing events in *Ahcy* and *Mat1a*. Exon usage for *Mat1a* and *Ahcy* transcripts at CT44, with exons plotted along genomic coordinates (left-to-right). Exons are numbered from the left-most coordinate regardless of which is the coding strand (*Ahcy* is coded on the – strand, meaning exon 1 is part of the 3’-UTR). The gene model integrating all known exons is shown under each graph, with known transcripts and their Ensembl IDs underneath. Exons coloured purple are significant (p <0.05) in DEXSeq at CT44.

**Figure S5:**
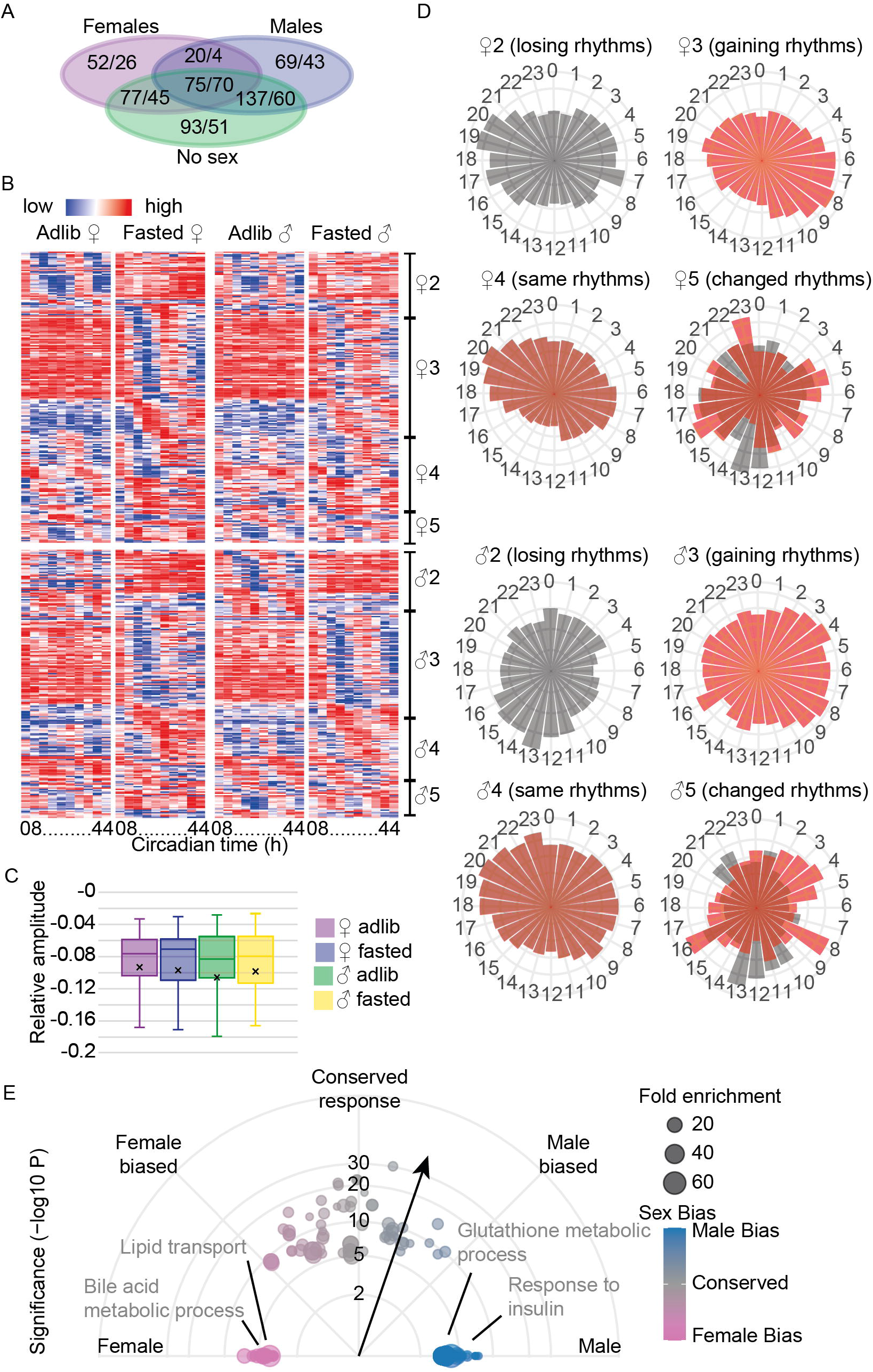
Sex-specific circadian responses of the hepatic alternative transcriptome. **A**, Venn diagram of differentially used exons/genes identified by dryR with female-only (N = 2), male-only (N = 2) or combined (N = 4) data. **B**, Heatmap of exons with altered rhythmicity (DEXSeq: q < 0.05, |log2FC| > 0.58 in ≥2 time points; dryR in ≥1 sex: BICW > 0.8), clustered by phase in *ad libitum* livers (models 2, 4 and 5) or fasted livers (model 3) in females (top heatmap) or male mice (bottom). Colours represent the mean of N = 2 males or 2 females; dryR performed independently with N = 2 per sex. **C**, Box-and-whisker plots of relative rhythmic amplitude (BICW > 0.8) per group; dots indicate means, horizontal lines indicate medians. Kruskal–Wallis test (p = 0.94). **D**, Polar plots of circadian phase differences between sexes for each model. **E**, Polar plot of aggregated GO results for exons in models 2, 3 and 5 in males or females. Vector length = −log10(FDR); dot size = fold enrichment; angle = sex bias (0°, female-specific; 90°, shared; 180°, male-specific). See also Supplemental Table S6.

**Supplemental Table S1:** Raw results of DESeq2 analyses performed for each time points, with diet as the main factor and sex as a covariate. Also includes unfiltered raw and normalised (median of ratio) counts. Available at the University of Manchester Figshare repository, https://doi.org/10.48420/32091148.

**Supplemental Table S2:** Raw results of maSigPro analyses with N = 4 mice or N = 2 males or females, accompanied with Enrichr and Gene Ontology analyses for genes in each maSigPro clusters that also meet DESeq2’s criteria, as described in the main text. Available at the University of Manchester Figshare repository, https://doi.org/10.48420/32091148.

**Supplemental Table S3:** Raw results of dryR analyses with N = 4 mice or N = 2 males or females, accompanied with Enrichr and Gene Ontology analyses with genes from model 2, 3 and 5 that also meet DESeq2’s criteria, as described in the main text. Available at the University of Manchester Figshare repository, https://doi.org/10.48420/32091148.

**Supplemental Table S4:** Raw results of DEXSeq analyses performed for each time points, with diet as the main factor and sex as a covariate. Available at the University of Manchester Figshare repository, https://doi.org/10.48420/32091148.

**Supplemental Table S5:** Raw results of maSigPro analyses on exon usage with N = 4 mice or N = 2 males or females, accompanied with Gene Ontology analyses with genes represented in the exons in each maSigPro clusters that also meet DEXSeq’s criteria, as described in the main text. Available at the University of Manchester Figshare repository, https://doi.org/10.48420/32091148.

**Supplemental Table S6:** Raw results of dryR analyses on exon usage with N = 4 mice or N = 2 males or females, accompanied with 4 Gene Ontology analyses: one with exons from model 2, 3 and 5 in males; one with exons from model 2, 3 and 5 in females; one “GO Circadian” with genes whose exons are rhythmic in adlib samples (model 2, 4 and 5 but not filtered by DEXSeq’s criteria); one “GO Fasting” with genes whose exons have been allocated to model 2, 3 and 5 in the dryR analysis on N = 4 mice. Available at the University of Manchester Figshare repository, https://doi.org/10.48420/32091148.

**Supplemental Table S7:** Raw and unfiltered exon counts, output of DEXSeq’s count function. Available at the University of Manchester Figshare repository, https://doi.org/10.48420/32091148.

## REFERENCES

1 Sokolovic, M. et al. The transcriptomic signature of fasting murine liver. BMC Genomics 9, 528, doi:10.1186/1471-2164-9-528 (2008).

2 Bauer, M. et al. Starvation response in mouse liver shows strong correlation with life-span-prolonging processes. Physiol Genomics 17, 230–244, doi:10.1152/physiolgenomics.00203.2003 (2004).

3 Schupp, M. et al. Metabolite and transcriptome analysis during fasting suggest a role for the p53-Ddit4 axis in major metabolic tissues. BMC Genomics 14, 758, doi:10.1186/1471-2164-14-758 (2013).

4 Zhang, F., Xu, X., Zhou, B., He, Z. & Zhai, Q. Gene expression profile change and associated physiological and pathological effects in mouse liver induced by fasting and refeeding. PLoS One 6, e27553, doi:10.1371/journal.pone.0027553 (2011).

5 Rennert, C. et al. The Diurnal Timing of Starvation Differently Impacts Murine Hepatic Gene Expression and Lipid Metabolism - A Systems Biology Analysis Using Self-Organizing Maps. Front Physiol 9, 1180, doi:10.3389/fphys.2018.01180 (2018).

6 Bake, T., Murphy, M., Morgan, D. G. & Mercer, J. G. Large, binge-type meals of high fat diet change feeding behaviour and entrain food anticipatory activity in mice. Appetite 77, 60–71, doi:10.1016/j.appet.2014.02.020 (2014).

7 Fonken, L. K. et al. Light at night increases body mass by shifting the time of food intake. Proc Natl Acad Sci U S A 107, 18664–18669, doi:10.1073/pnas.1008734107 (2010).

8 Vollmers, C. et al. Time of feeding and the intrinsic circadian clock drive rhythms in hepatic gene expression. Proc Natl Acad Sci U S A 106, 21453–21458, doi:10.1073/pnas.0909591106 (2009).

9 Kinouchi, K. et al. Fasting Imparts a Switch to Alternative Daily Pathways in Liver and Muscle. Cell Rep 25, 3299–3314 e3296, doi:10.1016/j.celrep.2018.11.077 (2018).

10 Della Torre, S. et al. Short-Term Fasting Reveals Amino Acid Metabolism as a Major Sex-Discriminating Factor in the Liver. Cell Metab 28, 256–267 e255, doi:10.1016/j.cmet.2018.05.021 (2018).

11 Wang, H. et al. Lipin 1 modulates mRNA splicing during fasting adaptation in liver. JCI Insight 6, doi:10.1172/jci.insight.150114 (2021).

12 McGlincy, N. J. et al. Regulation of alternative splicing by the circadian clock and food related cues. Genome Biol 13, R54, doi:10.1186/gb-2012-13-6-r54 (2012).

13 Nueda, M. J., Tarazona, S. & Conesa, A. Next maSigPro: updating maSigPro bioconductor package for RNA-seq time series. Bioinformatics 30, 2598–2602, doi:10.1093/bioinformatics/btu333 (2014).

14 Conesa, A., Nueda, M. J., Ferrer, A. & Talon, M. maSigPro: a method to identify significantly differential expression profiles in time-course microarray experiments. Bioinformatics 22, 1096–1102, doi:10.1093/bioinformatics/btl056 (2006).

15 Love, M. I., Huber, W. & Anders, S. Moderated estimation of fold change and dispersion for RNA-seq data with DESeq2. Genome Biol 15, 550, doi:10.1186/s13059-014-0550-8 (2014).

16 Thomas, P. D. et al. PANTHER: Making genome-scale phylogenetics accessible to all. Protein Sci 31, 8–22, doi:10.1002/pro.4218 (2022).

17 Mi, H. et al. Protocol Update for large-scale genome and gene function analysis with the PANTHER classification system (v.14.0). Nat Protoc 14, 703–721, doi:10.1038/s41596-019-0128-8 (2019).

18 Xie, Z. et al. Gene Set Knowledge Discovery with Enrichr. Curr Protoc 1, e90, doi:10.1002/cpz1.90 (2021).

19 Kuleshov, M. V. et al. Enrichr: a comprehensive gene set enrichment analysis web server 2016 update. Nucleic Acids Res 44, W90–97, doi:10.1093/nar/gkw377 (2016).

20 Chen, E. Y. et al. Enrichr: interactive and collaborative HTML5 gene list enrichment analysis tool. BMC Bioinformatics 14, 128, doi:10.1186/1471-2105-14-128 (2013).

21 Chen, M. et al. CD36 regulates diurnal glucose metabolism and hepatic clock to maintain glucose homeostasis in mice. iScience 26, 106524, doi:10.1016/j.isci.2023.106524 (2023).

22 Zheng, X., Yang, Z., Yue, Z., Alvarez, J. D. & Sehgal, A. FOXO and insulin signaling regulate sensitivity of the circadian clock to oxidative stress. Proc Natl Acad Sci U S A 104, 15899–15904, doi:10.1073/pnas.0701599104 (2007).

23 Qiu, S. et al. Hepatic estrogen receptor alpha is critical for regulation of gluconeogenesis and lipid metabolism in males. Sci Rep 7, 1661, doi:10.1038/s41598-017-01937-4 (2017).

24 Della Torre, S. et al. An Essential Role for Liver ERalpha in Coupling Hepatic Metabolism to the Reproductive Cycle. Cell Rep 15, 360–371, doi:10.1016/j.celrep.2016.03.019 (2016).

25 Damiola, F. et al. Restricted feeding uncouples circadian oscillators in peripheral tissues from the central pacemaker in the suprachiasmatic nucleus. Genes Dev 14, 2950–2961, doi:10.1101/gad.183500 (2000).

26 Kawamoto, T. et al. Effects of fasting and re-feeding on the expression of Dec1, Per1, and other clock-related genes. J Biochem 140, 401–408, doi:10.1093/jb/mvj165 (2006).

27 Sun, J. et al. Hepatocyte Period 1 dictates oxidative substrate selection independent of the core circadian clock. Cell Rep 43, 114865, doi:10.1016/j.celrep.2024.114865 (2024).

28 Weger, B. D. et al. Systematic analysis of differential rhythmic liver gene expression mediated by the circadian clock and feeding rhythms. Proc Natl Acad Sci U S A 118, doi:10.1073/pnas.2015803118 (2021).

29 Takahashi, J.S. Transcriptional architecture of the mammalian circadian clock. Nature reviews.Genetics 18, 164–179, doi:10.1007/978-3-319-27069-2_2 [doi] (2017).

30 Chen, L., Tovar-Corona, J. M. & Urrutia, A. O. Alternative splicing: a potential source of functional innovation in the eukaryotic genome. Int J Evol Biol 2012, 596274, doi:10.1155/2012/596274 (2012).

31 Anders, S., Reyes, A. & Huber, W. Detecting differential usage of exons from RNA-seq data. Genome Res 22, 2008–2017, doi:10.1101/gr.133744.111 (2012).

32 Saer, B., Taylor, G., Hayes, A., Ananthasubramaniam, B. & Fustin, J.-M. Dietary methionine/choline deficiency affects behavioural and molecular circadian rhythms. bioRxiv, 2024.2001.2004.574014, doi:10.1101/2024.01.04.574014 (2024).

33 Fukumoto, K. et al. Excess S-adenosylmethionine inhibits methylation via catabolism to adenine. Commun Biol 5, 313, doi:10.1038/s42003-022-03280-5 (2022).

34 Fustin, J. M. Methyl Metabolism and the Clock: An Ancient Story With New Perspectives. J Biol Rhythms, 7487304221083507, doi:10.1177/07487304221083507 (2022).

35 Fustin, J. M. et al. Methylation deficiency disrupts biological rhythms from bacteria to humans. Commun Biol 3, 211, doi:10.1038/s42003-020-0942-0 (2020).

36 Greco, C. M. et al. S-adenosyl-l-homocysteine hydrolase links methionine metabolism to the circadian clock and chromatin remodeling. Sci Adv 6, doi:10.1126/sciadv.abc5629 (2020).

37 Shima, H. et al. S-Adenosylmethionine Synthesis Is Regulated by Selective N(6)-Adenosine Methylation and mRNA Degradation Involving METTL16 and YTHDC1. Cell Rep 21, 3354–3363, doi:10.1016/j.celrep.2017.11.092 (2017).

38 Pendleton, K. E. et al. The U6 snRNA m(6)A Methyltransferase METTL16 Regulates SAM Synthetase Intron Retention. Cell 169, 824–835 e814, doi:10.1016/j.cell.2017.05.003 (2017).

39 Koike, N. et al. Transcriptional architecture and chromatin landscape of the core circadian clock in mammals. Science (New York, N.Y.) 338, 349–354, doi:10.1126/science.1226339 [doi] (2012).

40 Paschos, G. K., Lordan, R. & FitzGerald, G. A. Intersection of sex and circadian biology. Current Opinion in Physiology 45, 100834, doi:10.1016/j.cophys.2025.100834 (2025).

41 Astafev, A. A., Mezhnina, V., Poe, A., Jiang, P. & Kondratov, R. V. Sexual dimorphism of circadian liver transcriptome. iScience 27, 109483, doi:10.1016/j.isci.2024.109483 (2024).

42 Munyoki, S. K. et al. Intestinal microbial circadian rhythms drive sex differences in host immunity and metabolism. iScience 26, 107999, doi:10.1016/j.isci.2023.107999 (2023).

43 Bazhan, N. et al. Sex Differences in Liver, Adipose Tissue, and Muscle Transcriptional Response to Fasting and Refeeding in Mice. Cells 8, doi:10.3390/cells8121529 (2019).

44 Qian, J. et al. Sex differences in the circadian misalignment effects on energy regulation. Proc Natl Acad Sci U S A 116, 23806–23812, doi:10.1073/pnas.1914003116 (2019).

45 Goldstein, I. et al. Transcription factor assisted loading and enhancer dynamics dictate the hepatic fasting response. Genome Res 27, 427–439, doi:10.1101/gr.212175.116 (2017).

46 Fustin, J. M. et al. Two Ck1delta transcripts regulated by m6A methylation code for two antagonistic kinases in the control of the circadian clock. Proc Natl Acad Sci U S A 115, 5980–5985, doi:10.1073/pnas.1721371115 [doi] (2018).

47 Narasimamurthy, R. et al. CK1delta/epsilon protein kinase primes the PER2 circadian phosphoswitch. Proc Natl Acad Sci U S A 115, 5986–5991, doi:10.1073/pnas.1721076115 [doi] (2018).

48 Gene Ontology, C. The Gene Ontology knowledgebase in 2026. Nucleic Acids Res 54, D1779–D1792, doi:10.1093/nar/gkaf1292 (2026).

49 Ashburner, M. et al. Gene ontology: tool for the unification of biology. The Gene Ontology Consortium. Nat Genet 25, 25–29, doi:10.1038/75556 (2000).

50 Keenan, A. B. et al. ChEA3: transcription factor enrichment analysis by orthogonal omics integration. Nucleic Acids Res 47, W212–W224, doi:10.1093/nar/gkz446 (2019).

51 Lachmann, A. et al. ChEA: transcription factor regulation inferred from integrating genome-wide ChIP-X experiments. Bioinformatics 26, 2438–2444, doi:10.1093/bioinformatics/btq466 (2010).

